# Multi-Cell ECM compaction is predictable via superposition of nonlinear cell dynamics linearized in augmented state space

**DOI:** 10.1101/526426

**Authors:** Michaelle N Mayalu, Min-Cheol Kim, Harry Asada

## Abstract

Cells interacting through an extracellular matrix (ECM) exhibit emergent behaviors resulting from collective intercellular interaction. In wound healing and tissue development, characteristic compaction of ECM gel is induced by multiple cells that generate tensions in the ECM fibers and coordinate their actions with other cells. Computational prediction of collective cell-ECM interaction based on first principles is highly complex especially as the number of cells increase. Here, we introduce a computationally-efficient method for predicting nonlinear behaviors of multiple cells interacting mechanically through a 3-D ECM fiber network. The key enabling technique is superposition of single cell computational models to predict multicellular behaviors. While cell-ECM interactions are highly nonlinear, they can be linearized accurately with a unique method, termed Dual-Faceted Linearization. This method recasts the original nonlinear dynamics in an augmented space where the system behaves more linearly. The independent state variables are augmented by combining auxiliary variables that inform nonlinear elements involved in the system. This computational method involves a) expressing the original nonlinear state equations with two sets of linear dynamic equations b) reducing the order of the augmented linear system via principal component analysis and c) superposing individual single cell-ECM dynamics to predict collective behaviors of multiple cells. The method is computationally efficient compared to original nonlinear dynamic simulation and accurate compared to traditional Taylor expansion linearization. Furthermore, we reproduce reported experimental results of multi-cell induced ECM compaction.

**Author summary:** Collective behaviors of multiple cells interacting through an ECM are prohibitively complex to predict with a mechanistic computational model due to its highly nonlinear dynamics and high dimensional space. We introduce a methodology where nonlinear dynamics of single cells are superposed to predict collective multi-cellular behaviors through a developed linearization method. We represent nonlinear single cell dynamics with linear state equations by augmenting the independent state variables with a set of auxiliary variables. We then transform the linear augmented state equations to a low-dimensional latent model and superpose the linear latent models of individual cells to predict collective behaviors that emerge from multi-cellular interactions. The method successfully reproduced experimental results of cell-induced ECM compaction.

## Introduction

Cell-induced compaction of fibrous extracellular matrix (ECM) is an important mechanism for numerous processes such as wound healing and tissue development [1–3]. During wound healing, for example, traction forces exerted by fibroblasts and myofibroblasts result in ECM compaction at the site of injury [2, 3]. In vitro experiments using cell-populated collagen gel reveal global compaction of the matrix as a result of cooperative effect of multiple cells at the boundaries as well as propagation through the bulk [4–6]. Furthermore, matrix densification is observed in the regions around [7] and in-between cells. Here we examine the mechanical aspect of intercellular communication through the ECM and how contractile cells can induce emergent mechanical changes leading to matrix compaction. From a simplified mechanics point of view, compaction results when the traction forces exerted by the contractile cells embedded within the ECM overcome the resistive forces of the ECM structure, including viscoelastic forces and elastic energy forces. As a result the matrix is deformed from its original stress-free state and the elastic modulus increases [4–7].

In reality, the compaction process is far more complex. The ECM forms a network of cross-linked fibers that is highly nonlinear and intricate, but is critical for predicting large compaction and long-range transmission of forces [4]. As a large deformation is induced by contractile cells, the standard linear mechanics model yields substantial errors. The ECM fiber network is anisotropic and causes irreversible deformations as a large compaction takes place. This prominent nonlinearity prohibits use of simple methods for predicting the ECM compaction by a multitude of cells. In addition, cells can internally modulate their state in response to local mechanical stresses within the ECM, which influences cell polarity, contractility, stiffness and strength of focal adhesions [8, 9]. These cell properties are highly nonlinear and complex. Consideration of these nonlinear physical and physiological properties involved in the cell-ECM mechanics often result in differential equations that are intractably complex due to high-dimensional, nonlinear coupled dynamics.

Many in silico modeling approaches in the areas of wound healing and fibrotic disease have helped elucidate and explore the underlying phenomena involved in cell-induced ECM compaction, and have been used to supplement in vitro experiments for fast and inexpensive methods of evaluation. Approaches in previous works include: i) a hybrid continuum-discrete framework consisting of the macroscopic finite element domain and local microscopic fiber network [10], ii) rule-based models with deformable cells and ECM fibers to explain matrix remodeling and durotaxis [11, 12], iii) a discrete fiber model of cell populated fibrous matrix [13], and iv) continuum models of ECM gel compaction [7, 14, 15]. Even though these works provide many insights, they also simplify the ECM gel compaction mechanism by: a) 2-D representation of a 3-D system, b) exclusion of intracellular mechanics such as mechanobiology of actin stress fibers, focal adhesions, and remodeling of cellular and nuclear membranes, and c) consideration of linear elastic spring model of ECM fibers without including the viscoelastic nature of the fibers. Consequently, these prior models abstract detailed cell-ECM interactions, resulting in limitations to understanding how these interactions enable characteristic gel compaction.

In the current work, the ECM is modeled as a 3-D cross-linked network of discrete, viscoelastic fibers, and detailed mechanistic cell dynamics, including focal adhesion dynamics, cytoskeleton remodeling, actin motor activity and lamellipodia protrusion, are derived from basic principles. The resultant model is computationally complex, especially for a larger number of cells. The governing differential equations are highly nonlinear, coupled, and of high dimension. Here, we solved this difficulty by introducing a methodology having its disciplinary basis spanned in system dynamics, machine learning, and statistics.

It is known that a nonlinear system can behave more linearly when recast in a larger space [16]. In our approach, the original nonlinear dynamics derived from physical and physiological principles are recast in an enlarged state space by augmenting independent state variables with auxiliary variables that inform input-output characteristics of the nonlinear elements involved in the system. Once recast in the augmented space, the nonlinear system can be represented as an augmented set of linear dynamic equations. The linear representation facilitates model reduction using latent variable analysis, which can be shown is difficult to apply to highly nonlinear systems [17–20]. Furthermore, linearization in the augmented space allows for superposition of multiple subsystems. In the current work, collective behaviors of multiple cells are predicted via superposition of single cell subsystems through the linearization in the augmented state space. The proposed methodology is general, and is applicable to a broader class of problems where large-scale, collective behaviors must be predicted while retaining sufficient mechanistic details.

## Results

### Governing equations of collective cell behaviors in ECM fiber network

We construct a computational model for predicting cell-mediated gel compaction by multiple (*n_cell_*) cells having a uniform phenotype and interacting through a surrounding 3-D ECM fiber network. The ECM is modeled as a network of many fibers connected at a large number of nodes (*N_e_* ≈ 2000), whereas each cell is represented with a mesh structure consisting of multiple nodes (*N_c_* ≈ 200) which forms the cell outer membrane (see Fig-1A). The cell outer membrane deforms and gains traction as the nodes on the membrane bond to the nodes of the surrounding ECM fiber network and form focal adhesions, which occur when bonding molecules (or integrins) on the cell membrane bind to ligands on ECM.

**Fig 1.**
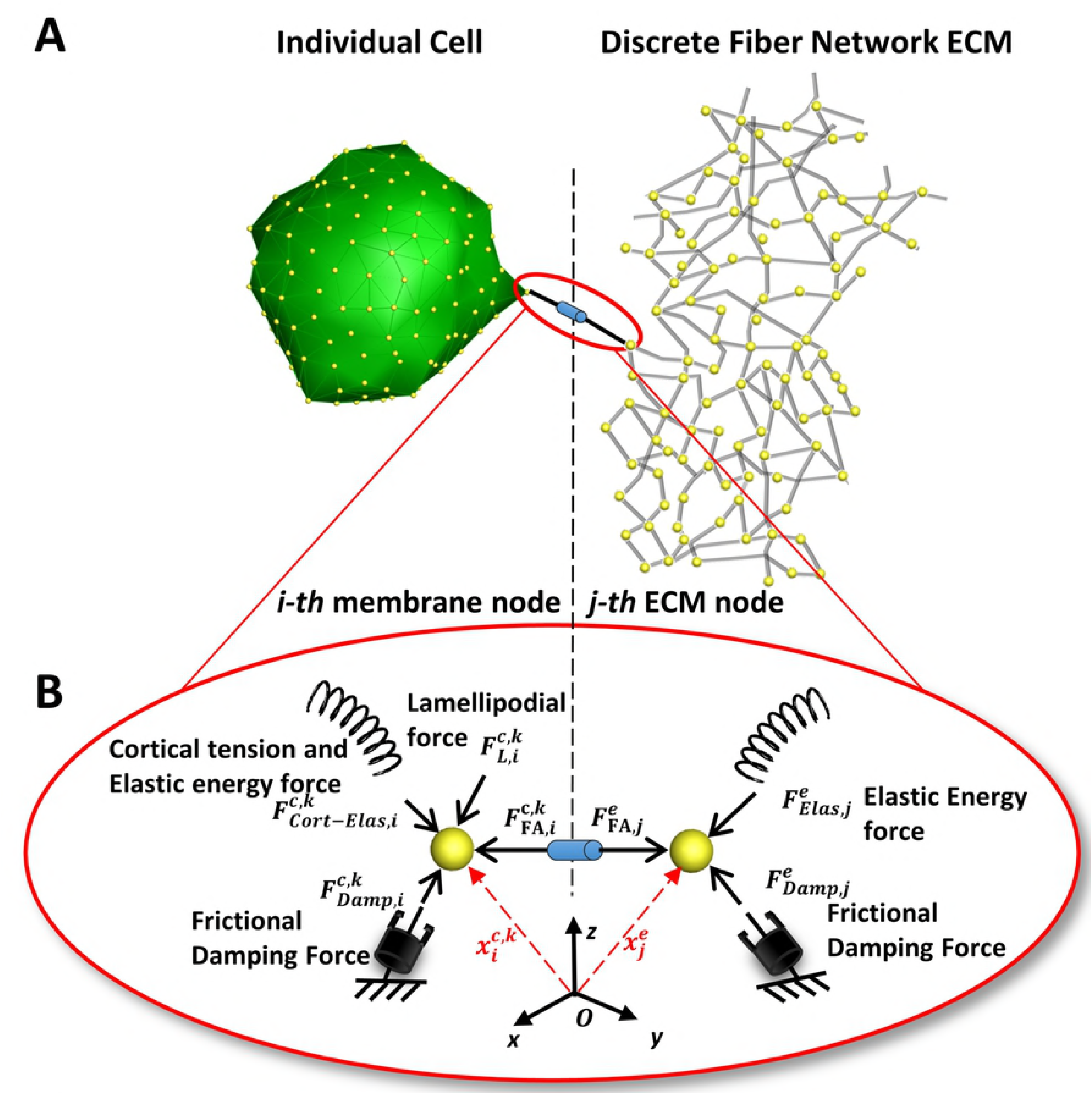
Schematic diagram of cell-ECM interaction. A: Each cell is represented by a mesh structure consisting of multiple nodes which are indicated by the yellow spheres. The ECM is modeled as a network of many fibers connected through a large number of nodes. The *i*-th membrane node is attached to the *j*-th ECM node through a focal adhesion connection. B: The forces acting on each membrane node include the cortical tension and membrane elastic energy force 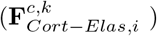, focal adhesion force 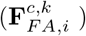, lamellipodium force 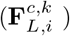, and frictional damping force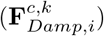. The forces acting on each node within the ECM fiber network include the elastic energy force 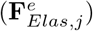, focal adhesion force 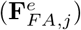, and damping force 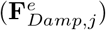.

Consider the *i*-th outer membrane node of the *k*-th cell with three dimensional spatial coordinates 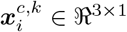 (See Fig-1B). The forces acting on it include the cell’s cortical tension force and elastic energy force (collectively denoted as 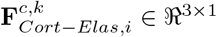), focal adhesion force (denoted as 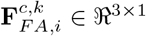), lamellipodium force 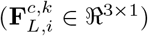, and frictional damping force 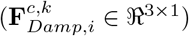 [21, 22]. Assuming that the mass of the node is negligibly small and the damping force is given by 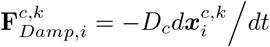, where *D_c_* is damping constant, the equation of motion is given by:

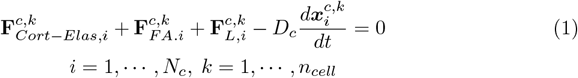

The generation of lamellipodium force pertains to the polarity of the cell. Namely, lamellipodia extend in a particular direction of the cell determined by the cell’s polarity [21–24]. The cell polarity and the lamellipodium forces can be treated as a cell’s decision or, in the system dynamics terminology, control inputs. Let 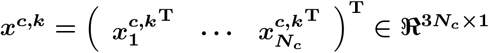 be a vector containing the 3-D coordinates of all the cell membrane nodes. Here the superscript in **X^T^** represents the transpose of matrix or vector **X**. The above equation of motion can be written collectively as:

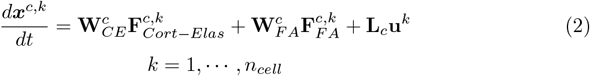

where 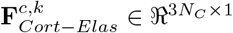 is a vector comprising cortical tension and elastic energy forces for all the cell nodes (*i* = 1, ⋯, *N_C_*), 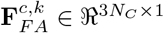 is a vector of focal adhesion forces at all the cell nodes, **u**^*k*^ ∈ ℜ^3*N*_*C*_×1^ is an input vector containing all the lamellipodium forces 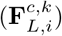, and 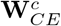 and **L**_*c*_ are constant matrices of consistent dimensions.

The equation of motion of the surrounding ECM fiber network can be represented in a similar manner. The forces acting on the *j*-th node of the fiber network are the elastic energy forces, including both lateral restoring forces and the one associated with bending moments, 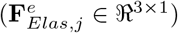, focal adhesion forces from the shared attachment with the cell 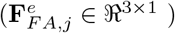 and damping forces 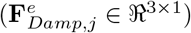 [21–24]. The equation of motion can be written as:

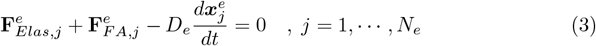

Let 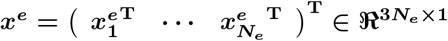 be a vector containing the 3-D coordinates of all the ECM nodes. Then equation 3 can be written as:

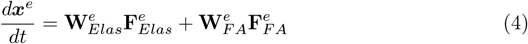

The ECM elastic energy force is a nonlinear function of ECM coordinates ***x**^e^*. The cortical tension and elastic energy force of the *k*-th cell is a nonlinear function of its membrane coordinates ***x**^c,k^*. Here ***x**^e^* and ***x**^c,k^* are independent state variables of the multi-cell ECM system.

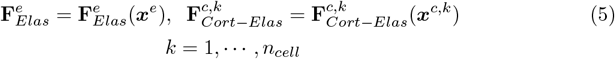

The focal adhesion force is modeled as a stochastic binding process between nodes on the cell membrane and those on the ECM. Using Monte Carlo simulations it has been found that focal adhesion forces can be approximated to a nonlinear algebraic function of cell membrane and ECM nodes as well as the biochemical parameters involved in integrin-ligand binding [21, 22].

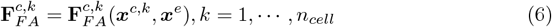

Assuming that no two cells bind to the same ECM node, we can find that the focal adhesion force of the *i*-th membrane node of the *k*-th cell attached to the *j*-th ECM node must satisfy:

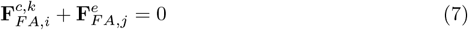

Namely, 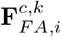 and 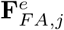 have the same magnitude with the opposite signs. Therefore, all the focal adhesion forces of the *k*-th cell can be mapped to the corresponding ECM nodes. Collectively, the focal adhesion forces of all the nodes within the ECM may be written as:

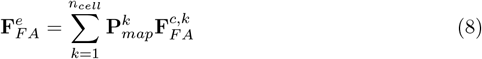

where 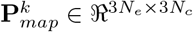 is a parameter matrix (consisting of either 0 or −1 elements) that maps the membrane focal adhesion forces of the individual cells 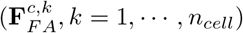 to the corresponding ECM focal adhesion forces 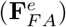. The focal adhesion connections between the membrane nodes and ECM nodes change over time as the cell membrane deforms, gains traction and generates lamellipodial protrusions. Therefore, the mapping matrix 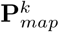 is updated at each time step. Details on the formation and structure of 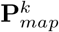 are given in the Methods Section. The *n_cell_* cells interact with each other through the surrounding ECM by generating focal adhesion forces, which propagate through the ECM fiber network and influence the other cells. The resultant collective behavior of the multiple cells is complex due to coupled, nonlinear dynamics.

### Dual-Faceted linearization

Although the governing equations derived above are rigorous and based on basic principles, they are complex and can become computationally expensive as the number of cells increases. Computational complexity is a key challenge in predicting collective behaviors of multiple cells. The number of state variables for the given system is 3*N_e_* + 3*N_c_n_cell_*, which is on the order of 7,000 for *n_cell_* = 2 and 9000 for *n_cell_* = 5. We aim to transform the governing equations into a linear latent variable representation in order to considerably reduce the number of state variables but also facilitate prediction of collective behaviors of the multiple cells through superposition of individual cell dynamics.

Model reduction is a challenging problem particularly for highly nonlinear, dynamical systems [19, 20, 25–27], as in the presented problem of collective behaviors of multiple cells within an ECM. If the system is linear or near linear, model reduction is more amenable and simple methods, such as Principal Component Analysis and Partial Least Squares, can reduce dimensionality. Here, we propose a unique linearization method, termed Dual-Faceted (DF) Linearization, and then apply a model reduction method to the linearized model. In DF Linearization, we represent the nonlinear dynamical system in an augmented space consisting of independent state variables (***x**^e^* and ***x**^c,k^*) and nonlinear forces 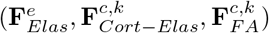 as the additional variables, termed auxiliary variables. Standard linearization, such as Taylor series expansion, is limited in accuracy, which may be valid only in the vicinity of a reference point. In DF Linearization, instead of taking “algebraic” linearization of these nonlinear terms, we consider “dynamic” linearization by representing their dynamic transitions using linear regressions.

Let the regression of the dynamic transition of auxiliary variable 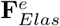, be expressed as:

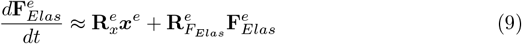

where 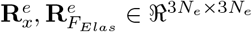 are parameter matrices. If an “algebraic” linearization using the Jacobian 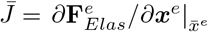 was utilized, the above equation would be: 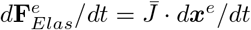.

This state transition equation through “algebraic” linearization is equivalent to one of the original independent state equation 4 because 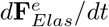 and *d**x**^e^*/*dt* are collinear within this formulation which renders it redundant. In contrast, the state transition equation presented in equation 9 is not collinear, providing a diverse facet of the nonlinear system. Similarly, for the auxiliary variables 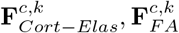, let the regression equations be written as:

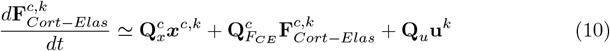

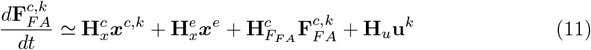

where 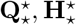 (⋆-corresponding subscript and superscript) are parameter matrices with consistent dimensions. The high-dimensional parameter matrices 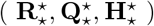 do not need to be determined explicitly as discussed in the subsequent sections. DF Linearization represents a nonlinear dynamical system with two sets of differential equations. One set is the original state equations governing the transition of the independent state variables and the other set is the regression of the dynamics of auxiliary variables. The original state equations, 2 and 4, are apparently linear in terms of the auxiliary variables and inputs. In these equations, all the forces acting on each node sum to zero. These are linear expressions when the nonlinear forces are treated as auxiliary variables. In addition, the auxiliary state transitions (equations 9 – 11) are given by linear regressions in the augmented space. Therefore, both differential equations are linear. The two linear differential equations represent different (or dual) facets of the original nonlinear system viewed from the augmented space, thus providing a richer representation of the nonlinearity.

### Latent variable transformation and model order reduction

Now that the original nonlinear system has been represented as a linear dynamical system in the augmented space, we can apply a latent variable modeling method to reduce model order. Represented in the augmented space, the differential equations may contain similar modes, or some variables are close to collinear. These similar modes and collinear variables can be eliminated by model order reduction methods.

Let *ζ^c,k^* be the augmented variable vector containing membrane node coordinates and forces of the *k*-th cell.

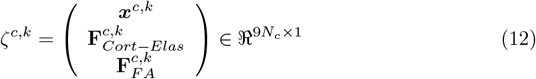

Here **u**^*k*^ (the cell’s lamellipodial force) is treated as input variables that are excluded from the augmented state space. Similarly, let *ζ^e^* be the augmented variable vector containing ECM node coordinates and forces:

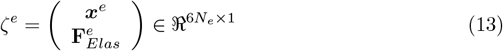

Focal adhesion forces 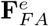 are determined by the individual cells by equation 8 and, thereby, excluded from the augmented space of the ECM.

We apply latent space analysis to vectors *ζ^c,k^* and *ζ^e^*, respectively. First we generate data by using equation 2, and 4 – 7. Computation of the nonlinear state equations is amenable for a single cell interacting with ECM. A data set can be created by simulating those nonlinear equations by placing a single cell at diverse locations, i.e. repeating the simulation with different initial conditions. Let 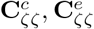 be the covariance matrices of simulation data sets of augmented states *ζ^c,k^* and *ζ^e^*, respectively. Each covariance matrix contains both independent state and auxiliary variables, where the latter is nonlinear functions of the former. If auxuliary variables were linear functions of the state variables, then the rank of the covariance matrix would be equal to the number of independent state variables. However due to the nonlinearity, the rank is higher. Details on the formation of 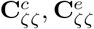 are given in the Methods Section.

The covariance analysis also reveals that the system represented in the augmented space contains many components that may be negligibly small. Using Principal Component Analysis, the original data of *ζ^c,k^* and *ζ^e^* can be represented with latent variables of truncated dimension *m_c_* ≪ 9*N_c_* and *m_e_* ≪ 6*N_e_*, respectively [27]:

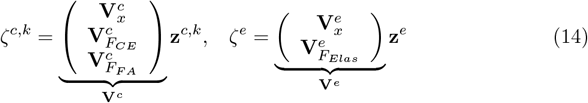

where matrices **V**^*c*^ ∈ ℜ^9*N_c_*×*m*^ and **V**^*e*^ ∈ ℜ^6*N_e_*×*m*^ are orthogonal matrices comprising the eigenvectors of the covariance matrices, and

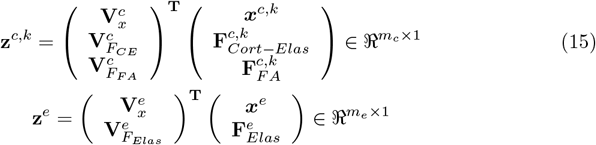

Differentiating the latent variable state vector **z^c,k^** and substituting equations 2, 10, 11 and equation 14 yields:

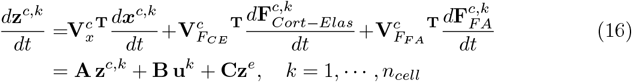

where:

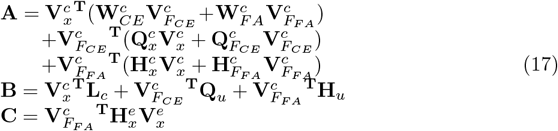

equation 16 provides the latent variable state equation of the *k*-th cell interacting with the ECM. Given the latent variable state of ECM **z**^*e*^ and the input **u**^*k*^ reflecting the cell’s decision, the transition of the cell’s latent variable state is determined locally without directly including the states of the other cells. Cells interact indirectly through the strain field created by other cells over the ECM fiber network.

The ECM dynamics can be represented in the latent variable space spanned by **V^e^**. Differentiating the latent variable state vector and substituting equations 4, 9 and 14 yield:

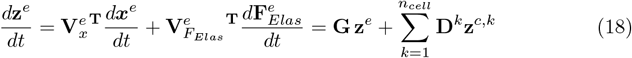

where:

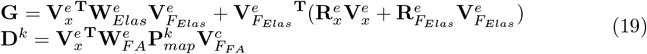

Fig-2 shows numerical examples of the DF linearization and subsequent latent variable transformation in reproducing accurate cell morphologies of the original nonlinear computational model over time. Remarkably, the linearized latent variable model can correctly reconstruct the complex cell membrane topology with *m* = *m_c_* + *m_e_* = 50 + 50 = 100 total latent variables. Fig-2B quantifies the root mean square error and computation time as a function of latent variable model dimension. As can be seen, the computation time for the latent variable cell-ECM system increases with increased latent variables while the root mean square error decreases. Conversely, the standard Taylor expansion linearization is not capable of representing cell morphologies without marked error which is quantified in Fig-2C. These results demonstrate the effectiveness of DF Linearization and model reduction in reconstructing simulations from a high dimensional complex nonlinear model.

**Fig 2.**
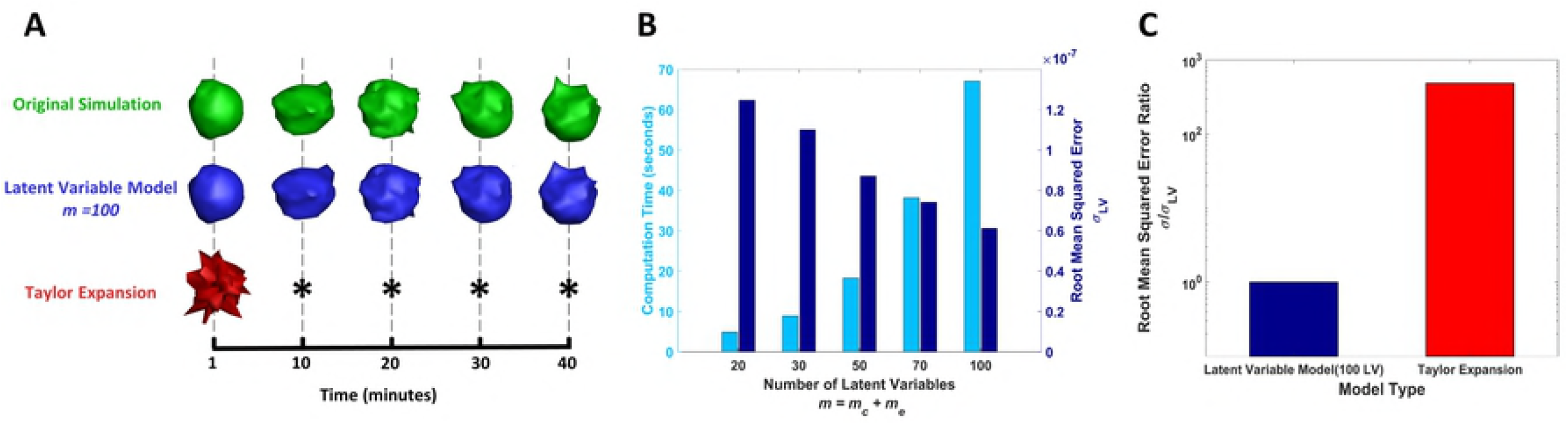
Comparison of nonlinear computational model, latent variable model, and taylor series expansion model for predicting a cell interacting with ECM. A: The cell morphologies over time for the original full nonlinear simulation (green), the latent variable simulation using 100 latent variables (blue), and the Taylor series expansion model (red). The Taylor series expansion model is not capable of representing cell morphologies without significant error (represented by *). B: The computation time and root mean squared error (*σ_LV_*) of the latent variable model as a function of the number of latent variables used to create the model. C: Comparison of root mean squared error of latent variable model using 100 latent variables and a model created by first order Taylor expansion of nonlinear terms.

This latent space model provides not only a low-dimensional structure for efficient computation, but also contains natural insights into the interactions among the multiple cells. Fig-3A shows the dynamic interactions in block diagram form based on equation 16 and equation 18. The ECM changes its latent variable state **z**^*e*^ with the autoregressive feedback through matrix **G** as well as with the forward path that collects the latent variable states of all the individual cells, as shown by the summing junction Σ. Each single cell changes its latent variable state **z**^*c,k*^ (*k* = 1,…, *n_cell_*) with autoregressive feedback through **A** as well as with two forward paths. The first path (fed through **C**) represents global feedback from the ECM (**z**^*e*^). The second path (fed through **B**) represents the updated lamellipodial forces **u**^*k*^ determined from ECM state **z**^*e*^. The lamellipodial forces can be thought of as the individual cell’s decision based on its position and updated ECM properties as explained more in the following section. The actions taken by all the cells are integrated into the global ECM state transition, which is fed back to the individual cells. Therefore, each cell is connected to other cells through the global feedback of the ECM latent variable state **z**^*e*^. Fig-3A manifests the control-theoretic interpretation of multiple cells interacting through ECM.

**Fig 3.**
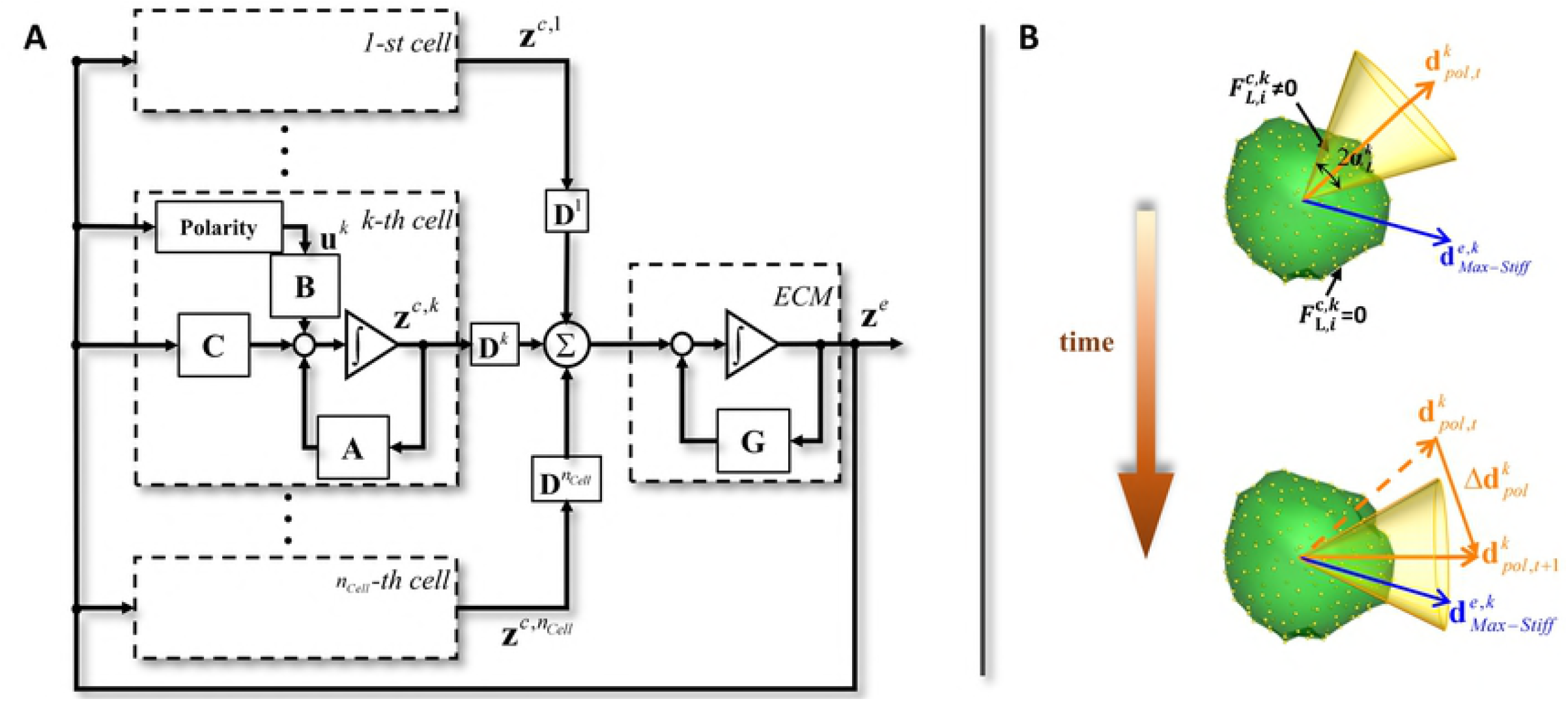
Block diagram of latent variable superposition model and schematic of the relationship between polarity direction, leading Eege and direction of maximum stiffness. A: The ECM changes its latent state with the autoregressive feedback through matrix G as well as with the feedforward path which collects the latent variable states of all the individual cells **z**^*c,k*^(*k* = 1,…, *n_cell_*) through matrices **D**^*k*^. Each cell changes its latent variable state with autoregressive feedback through **A** and are exposed to the ECM forces represented by latent vector **z**^*e*^ in two separate paths. The path through the cell polarity block and matrix B can be viewed as an “active input”. This feedback path includes a cell’s internal decision as to which direction it extends lamellipodia. In contrast, the other feedback path through a gain matrix **C** does not have a high-level cell decision, but is reactive, playing a “passive role”. B: The cell polarity direction rotates dynamically in such a way that the polarity vector 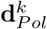 may align with the direction of the maximum stiffness 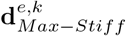. The leading edge of the cell is indicated by a right circular cone with apex angle 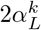 having its centerline aligned with the polarity direction. The membrane nodes of the *k*-th cell within the cone have nonzero lamellipodial forces 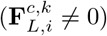. Membrane nodes outside this cone have zero lamellipodial forces 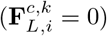.

Since the system is represented in a lower dimensional space, the high dimensional regression coefficient matrices 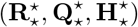 are not computed explicitly. Instead, the lower dimension coefficient matrices **A, B, C, G** are computed from numerical simulation data that can be transformed into latent variable space. Details are given in the Methods Section.

### Polarity and cell decisions

As discussed previously, input vector **u**^*k*^ pertains to the lamellipodial forces generated at each membrane node within the leading edge of the cell. The cell continuously updates its lamellipodial protrusions depending on the orientation of the leading edge as the cell’s polarization (or polarity) changes. The polarity of a cell is important to determine the orientation of the leading edge and is influenced by the direction of local maximum stiffness in the ECM [21–24]. Here we aim to extend the dynamics model of cell polarity developed in [21–23] to predict the formation of lamellipodia. Let 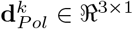 be a 3-dimensional unit vector indicating the direction of polarity in the *k*-th cell and 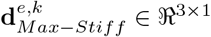 be a 3-dimensional unit vector pointing in the direction of the maximum stiffness of ECM in the vicinity of the *k*-th cell’s current location. According to [23], the cell polarity rotates dynamically in response to ECM’s local stiffness in such a way that the polarity vector may align with the direction of the maximum stiffness:

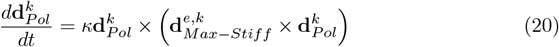

where × indicates vector product, and *κ* is a scalar parameter. Fig-3B illustrates this relationship. The polarity vector 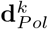 tends to align with the maximum stiffness direction, 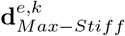.

The leading edge of the cell is indicated by a right circular cone with apex angle 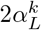 having its centerline aligned with the polarity direction. The membrane nodes of the *k*-th cell within the cone have nonzero lamellipodial forces 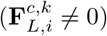. Membrane nodes outside this cone have zero lamellipodial forces 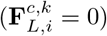. The direction of maximum ECM stiffness 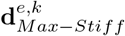 depends on the stress field within the ECM, which pertains to the latent state vector of ECM **z**^*e*^. The details are given in S3 Appendix.

### Computational analysis of ECM compaction by multiple cells

Compaction of the ECM by the collective efforts of multiple cells is numerically analyzed based on the model reduction and superposition of the nonlinear cell-ECM dynamics via DF Linearization. We first consider the case where two cells placed 30 *μ*m apart are embedded in a 3-D cylindrical ECM that measures 40 *μ*m in diameter and 100 *μ*m in length as seen in Fig-4A. The boundary conditions of the ECM fiber network are set such that the two flat planes on sides are fixed to space (constrained), while the curved surface surrounding the ECM is kept free (unconstrained). The volume of the cylindrical ECM shrinks over time from its initial unstressed state as the cells interact with the surrounding ECM. To quantify the spatiotemporal compaction process, the original ECM cylinder is segmented into 10 slices of 10 *μ*m thickness along its longitudinal axis as shown in Fig-4B. The volumetric changes to the individual slices are plotted in Fig-4C. The prediction of decreased cell volume by the latent variable superposition model (blue) agrees well with the ground-truth, full-scale nonlinear simulation results (green). This is further verified by the corresponding cross-sectional images of the 2-cell cylindrical ECM simulations in Fig-4C. The polarity directions of both cells (shown by red arrows initially pointing in arbitrary directions) shift to point inward, indicating that larger stresses are detected in the area between the cells. A video of the simulation comparison is shown in S1 Video.

**Fig 4.**
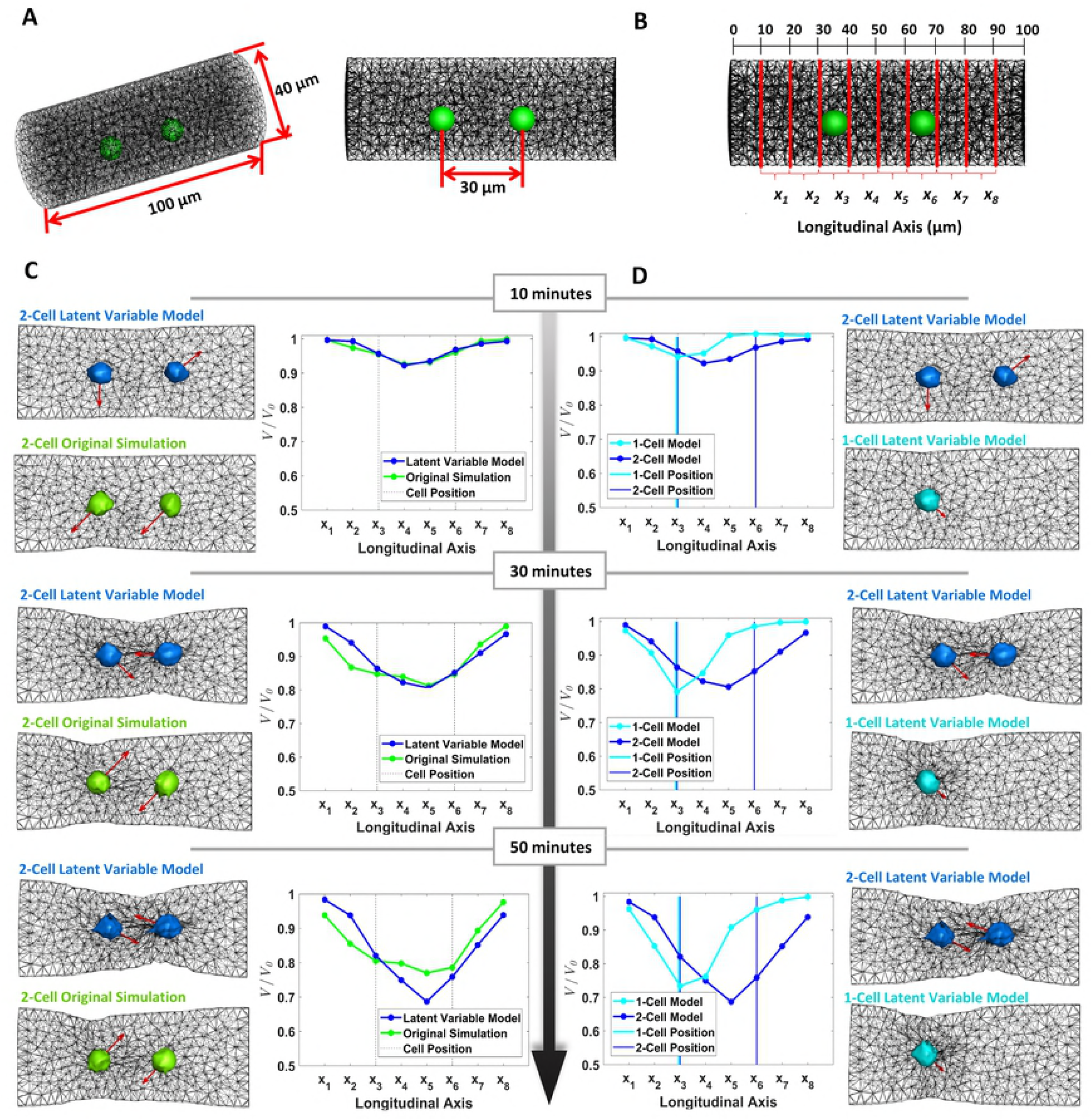
Comparison of ECM compaction between nonlinear computational model and linear latent variable models. A: Two cells placed 30 *μ*m apart are embedded in a 3-D cylindrical ECM that measures 40 *μ*m in diameter and 100 *μ*m in length. B: The ECM is subdivided along the longitudinal axis into 10 *μ*m length cylinders. The volume (V) of each subdivided cylinder may be estimated at multiple time points during the compaction simulation. C: ECM comparison at the subdivided segments at time *t* = 10 min, 30 min, and 50 min. Compaction predicted by the latent variable model simulations (blue) agrees well with the full nonlinear simulations (green). The compaction volume is normalized with the initial volume of each segment. The compaction is most significant in-between the cells (the region between the dashed lines in the plots). This is further verified by the corresponding cross-sectional images of the 2-cell cylindrical ECM simulations. Polarity directions of both cells (red arrows initially pointing in arbitrary directions) shift to point inward, indicating that larger stresses are detected in the area between the cells. D: Comparison of single cell (cyan) and two cell (blue) compaction predicted by the latent variable superposition model. As can be seen, the single cell model predicts more localized shrinkage of the ECM volume from its original unstressed state whereas the two cell model shows more global shrinkage extended to within the region between cells.

The proposed model is able to reproduce collective behaviors of multiple cells causing the characteristic compaction of ECM gel, which is not observed for single isolated cell models. This is further verified in Fig-4D which compares the ECM compaction results between single cell and two cell models. As can be seen, the single cell model predicts more localized shrinkage of the ECM volume whereas the two cell model shows more global shrinkage extended to within the region between the cells. A video of the simulation comparison is shown in S2 Video.

Fig-4D suggests the presence of more than one cell is necessary for the pronounced ECM compaction leading to emergent changes within the ECM. However, the emergence of pronounced compaction entails not only plurality of cells but proper cell spacing. Fig-5A shows that, as the spacing between cells increases, compaction is less pronounced between them, indicating decreased interaction and integration of cell induced propagated forces. This is summarized in Fig-5B which quantifies the average ECM elastic force in-between cells against cell spacing. From Fig-5B, we see that the average ECM elastic force in-between cells spaced at 100 *μ*m is an order of magnitude less than that of the cells spaced at 30 *μ*m. A video of this simulation is shown in S3 Video.

**Fig 5.**
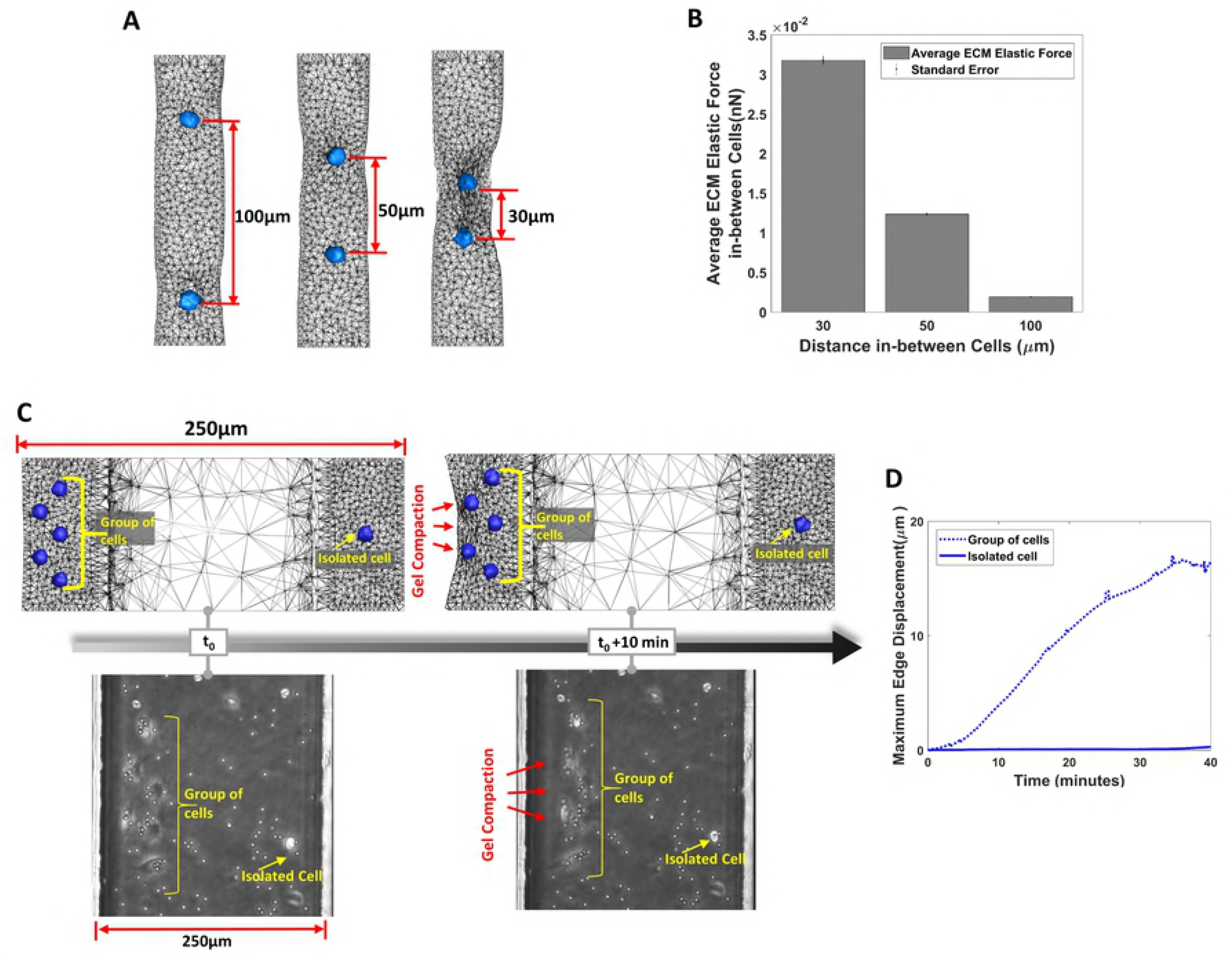
Effect of cell spacing and density on ECM compaction. (A) ECM compaction by two cells embedded within a cylindrical ECM spaced at 100*μm*, 50*μm*, and 30 *μm*. As the spacing between cells increases, compaction is less pronounced between them. (B) The average ECM elastic force in-between cells spaced at 100*μm* is an order of magnitude less than the elastic force in-between cells spaced at 30*μm*. (C) The latent variable superposition simulation (upper) can be used to reflect the behavior of heterogeneous planar distribution of MC3T3-E1 osteoblasts (lower) [Adapted from supplementary Material in [5], with permission of The Royal Society of Chemistry. Copyright 2009]. Whereas the group of cells at the left edge contract the gel, the isolated cell at the right edge did not contract the gel, indicating the importance of cell density for compaction. (D) For the latent variable superposition simulation the maximum displacement along the ECM edge noticeably increases over time for the group of cells and is minimal for the isolated cell.

The above computational results shown in Fig-4 and Fig-5 verify that the proposed method can capture collective behaviors of multiple cells. The verification was made by comparing the reduced-order superposition model using DF Linearization against the full-scale, nonlinear model. To further verify the capability of the reduced-order superposition model, a comparison is also made against in vitro experimental data of ECM compaction by a larger number of cells. As shown in Fig-5C and D, the computational model successfully reproduces the in vitro experiment conducted by Fernandez, et al [5], in which a heterogeneous planar distribution of MC3T3-E1 osteoblasts were plated in 3-D rectangular prism collagen gel of 50 *μ*m height, 100*μ*m width and 250 *μ*m length. The boundary conditions of ECM in the computational model were set to be consistent with experimental conditions in [5].

The multi-cell latent variable simulation is able to predict the characteristics of the ECM over time. Whereas the group of 5 cells at the left edge exhibit anisotropic contraction of the ECM at the boundary, the single isolated cell at the right edge barely contracts the gel. The experimental image (adapted from supplementary material in [5], with permission of The Royal Society of Chemistry. Copyright 2009) is also shown for comparison. Fig-5C compares the isolated cell to the group of cells in terms of maximum edge displacement. The isolated cell’s ECM edge displacement is so small that five times its displacement is still substantially lower than the displacement of the group of cells. A video of this simulation is shown in the S4 Video. The presented method for predicting collective behaviors of cell-mediated ECM gel compaction is scalable. Since the individual cell-ECM interactions are local computations, as given by equation 16, the computational complexity does not increase exponentially, although the number of cells increases.

## Methods

Implementation of the presented method is enabled by four key constructs: 1) generation of simulated training data, 2) formation of the data covariance matrix necessary for latent variable transformations, 3) estimation of the parameter matrices involved in the latent variable space state equations (Eq. 16 and Eq. 18), and 4) association of cell and ECM focal adhesion force variables using mapping matrix. Each method is detailed below.

### Generation of simulated training data based on the original nonlinear equations of single cells

The nonlinear state equations equation 2, 4–6 were computed with custom C-code based on references [21, 22]. The computation of a single isolated cell embedded in the cylindrical ECM took approximately 24 hours for a single simulation of physical time *T* ≈ 3600 seconds with sampling interval of 1 second. The simulation was repeated for *N* ≈ 10 times at different initial cell locations each time. The simulations with a single isolated cell embedded in the large rectangular ECM for reproducing the experimental result were run for approximately 5 days (120 hours). The physical time of simulation was *T* ≈ 3600. The simulation was repeated for *N* ≈ 10 for various initial locations of a cell. For each simulation, the total number of sample points for all the variables was over 5,000,000. The number of sample points was 5 × 10^7^. The computation was performed on Intel Xeon CPU E5-2687W @ 3.10 GHz (2 processors) with 32 logical cores. More details on the formulation of the full-scale nonlinear state equations are summarized in S1 Appendix.

### Formation of the data covariance matrix necessary for latent variable transformations

We create the covariance matrix using simulated training data. The training data consists of 3,600 time points of both state and auxiliary variables of a cell embedded in an ECM environment. The simulation is repeated *N* =10 times, each time with the cell embedded in distinct locations within the ECM. Then the data covariance matrices may be formed:

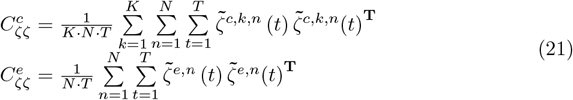

where 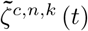 represents the mean centered t-th time sample (of augmented variable vector *ζ^c,k^*) for the k-th cell (here *K* = 1 or 2) embedded within for the ECM at the n-th simulation, and 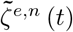 represents the mean centered t-th time sample of the augmented variable vector *ζ_e_* in the n-th simulation. By performing eigen-decomposition on the covariance matrices we obtain the orthogonal matrix **V**^*c*^, **V**^*e*^ comprised the eigenvectors of the data covariance matrix:

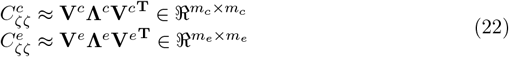

where **Λ**^*c*^, **Λ**^*e*^ are diagonal matrices containing the largest *m_c_* and *m_e_* eigenvalues of the covariance matrices, respectively. It is important to check whether the covariance matrices contain sufficiently rich data, and their first *m_c_* and *m_e_* components are sufficient to capture the cell-ECM dynamics at any cell location within the ECM. Standard techniques can be applied to validate the data and truncation of components [27]. With these, the ECM dynamics of a cell embedded within the ECM at an arbitrary location will be well represented in linear latent variable space which is critical to the success of the method. Covariance matrix calculation were conducted by using Matlab.

### Estimation of parameter matrices involved in latent variable space state equations

The following outlines the steps to compute coefficient matrices **A, B, C, G** by equation 16 and equation 18:

1. Create training data by simulating the original state equations, 2 and 4, using the full-scale, nonlinear model of single cells, as described above.
2. Compute covariance matrices 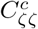, 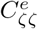, and obtain eigenvalues and eigenvectors **V**^*c*^ and **V**^*e*^, as described in the Methods Section.
3. Transform the data of the augmented state variables to latent variable space (**z**^*c,k,n*^ (*t*) and **z**^*e,n*^(*t*)) using the orthogonal matrices **V**^*c*^ and **V**^*e*^.
4. Compute time derivatives *d***z**^*c,k,n*^/*dt* and *d***z**^*e,n*^/*dt*, using latent variable space time samples and form a dataset:

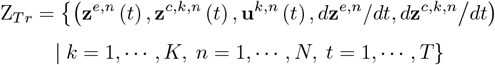

identify the parameter matrices A, B, C, G involved in the latent variable space state equations using Least Squares Estimate. The details about the Least Squares Estimate computation are given in S2 Appendix.

Parameter matrices **A** ∈ ℜ^*m_c_*×*m_c_*^, **B** ∈ ℜ^*m_c_*×3*N_c_*^, **C** ∈ ℜ^*m_c_*×*m_e_*^, **G** ∈ ℜ^*m_e_*×*m_e_*^ are substantially lower in dimension than the regression coefficient matrices 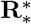, 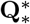, 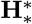 given in equation 9, equation 10 and equation 11. Therefore, fewer data points allow us to determine these parameter matrices in the latent variable space. It should be noted that matrices **D**^*k*^’s are of high dimension, but are not computed with regression since they consist of known matrices as shown in equation 19. Matlab was used for estimation of parameter matrices and subsequent computations of the latent variable model. 3-D visualization of simulation data was conducted using Tecplot 360.

### Association of cell and ECM focal adhesion force variables using mapping matrix

When a focal adhesion is formed between the i-th node of the k-th cell and the j-th node of ECM, the two focal adhesion forces sum to zero, as described previously (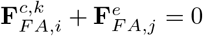 where 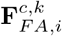, 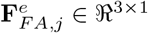). Representing this relationship in terms of the collective focal adhesion force vectors, 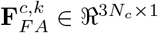 and 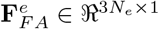, requires a matrix 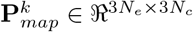. Let 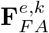 be the forces acting on the ECM nodes caused by focal adhesion between the k-th cell and the ECM nodes. This can be written

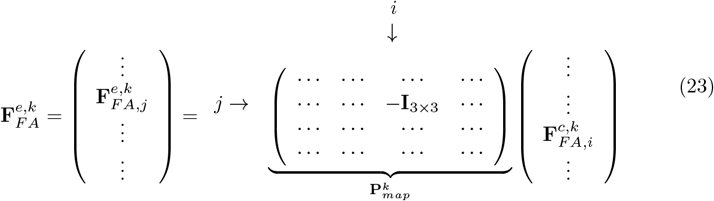

where **I**_3×3_ is the 3-dimensional identity matrix. Obtaining this mapping matrix 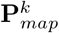 for all the cells, the complete focal adhesion forces in the ECM can be expressed in relation to the cell’s focal adhesion forces.

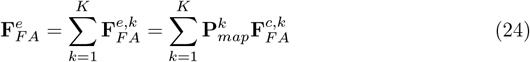

As previously mentioned, the focal adhesion connections between the cell membrane nodes and ECM nodes can vary over time as the cell membrane deforms, gains traction, and generates lamellipodial protrusions. Therefore, 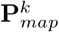 is updated to reflect the new focal adhesion attachments and detachments at each time step. The original nonlinear computational model has developed a functional relationship between the focal adhesion force, number of integrins, and distance between the membrane and ECM node (see details supplementary materials S1 Appendix). In the presented framework, the change in focal adhesion attachments can be derived from simulated training of the nonlinear computational model.

## Discussion

The collective ECM compaction by multiple cells is predicted through superposition of individual cells’ contributions in latent variable space. This is made possible by DF Linearization, latent variable transformation and subsequent superposition of single-cell models to predict the collective behavior among multiple cells.

As shown in Fig-3A, the DF Linearization has two-order-of-magnitude higher accuracy than the first-order Taylor expansion, and can approximate the original full scale model with a reasonable root-mean-square error. This representation of nonlinear dynamics is markedly different from standard linearization methods. To better understand the mechanism of DF Linearization, consider a simple example: a system consisting of one spring and a damping element with negligibly small mass, 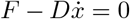. If the spring is a linear spring, *F* = *kx*, there is absolutely no difference between the equation in terms of state variable *x*, 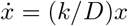, and the one in force 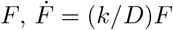. However, it is not the case if the spring is nonlinear, for example a hard spring: *F* = *ax* + *bx*^3^ where *a* > 0, *b* > 0. Representing the differential equation in two variables, one with the state variable *x* and the other with the auxiliary variable *F*, provide different equations.

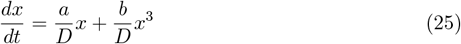

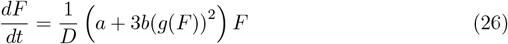

where *x* = *g*(*F*) is the inverse function of *F* = *ax* + *bx*^3^. Both equations represent the same nonlinear system, yet the expressions are different, hence Dual-Faceted representations. Linearizing these differential equations lead to two linear differential equations viewed from the augmented space. Note that equation 25 can be represented as a linear equation by using both state and auxiliary variables:

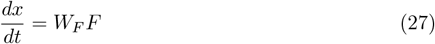

where *W_F_* = 1/*D*. The augmented state equation 26 can be approximated to a linear regression:

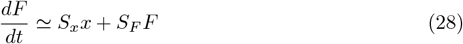

where *S_x_, S_F_* are regression parameters.

The expression given by equation 28 differs from the one based on the first order Taylor expansion (or “algebraic” linearization) which yields:

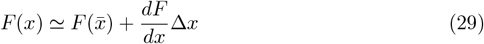

Furthermore, if we evaluate the derivative *J*(*x*) ≡ *dF*/*dx* at a particular point, 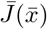, and then use equation 29 to express the augmented state equation, it reduces to 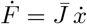. This implies that 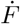 and 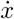 are proportional. Using an “algebraic” linearization yields a differential equation representing the transition of *F* that is collinear to the one representing the transition of *x*, and thereby an auxiliary state equation would not provide any new information.

Conversely, the regression model in equation 28 provides us with a diverse view of the original nonlinear system, thus providing a richer representation of the nonlinearity than the standard first order Taylor expansion. The traditional Taylor expansion method can deviate from the correct trajectory in a small number of iterations. However, the DF Linearization can track the true trajectories of both independent state and auxiliary variables with higher accuracy. The DF Linearization can predict accurate dynamic responses for limited yet sufficient intervals.

As applied to the analysis of multi-cell ECM compaction, linear augmented equations describing single cell-ECM interactions were derived from DF linearization, and then converted to a reduced-order linear representation by transformation onto a basis of eigenvectors derived from simulated data set. Unlike model reduction of nonlinear dynamical systems, which still remains a challenging problem in the field [17–20], the model reduction of a linear system through DF Linearization is straightforward. It allows for the evolution of independent and auxiliary states to be described within a lower dimensional linear manifold. The resulting reduced order latent variable model is capable of reproducing nonlinear dynamics, and the linearized structure of individual models facilitated their integration to describe multi-cell behaviors. The prediction of collective behaviors of a group of cells was achieved by superposing contributions of individual cells represented by latent variables **z**^*c,k*^, which evolves based on their own dynamics in response to the global ECM state represented by latent variable **z**^*e*^.

The linear representation of the collective multi-cell-ECM interactions manifests the two types of feedback actions by the individual cells. As shown in the block diagram in Fig-3A, the individual cells are exposed to the ECM forces represented by latent variable vector **z**^*e*^ in two separate paths. The path through the cell polarity block and matrix **B**, leading to lamellipodia formation, can be viewed as an “active input” as addressed in [5]. This feedback path includes a cell’s internal decision as to which direction it extends lamellipodia. In contrast, the other feedback path through a gain matrix **C** does not have a high-level cell decision, but is reactive, playing a “passive role” [5]. These feedback interactions support the prior experimental work [5]. It is interesting to note that ECM compaction begins almost instantaneously, but the magnitude of compaction is rather limited. Once the “active” feedback loop is initiated in, the ECM compacts further, resulting in a large deformation. As the polarity dynamics are rather slow, the second stage ECM compaction does not start immediately. The time scale is determined by the constant *κ* involved in the polarity dynamics equation 20. Using the proposed methodologies, we are able to reproduce intercellular mechanical interactions consistent with published experimental observations. In particular, the global compaction of gel volume via collective cell-contractile activities is characteristically different from local deformations of single isolated cells embedded within the same gel. Through study of emergent behaviors of groups of cells embedded in a 3-D ECM fiber network, we can advance our understanding of intercellular mechanical signaling during tissue formation [1–7]. There are a few limitations to our method, however. While the presented method can predict complex nonlinear behaviors, the method is still a type of approximation. Care must be taken with the validity period. In Fig-4C at the sample time of t = 50 minutes, the latent variable superposition simulation over predicts the volume shrinkage by 12%. With the current mathematical formulation, we have not yet incorporated the degradation of ECM fibers through matrix metalloproteinases. ECM degradation would be necessary to reproduce sustained movement and migration of the cells particularly in 3-D embedded matrices [28]. Since ECM degradation continuously changes the fiber connectivity through ECM remodeling, a methodology to update the node grid structure describing the ECM field would need to be developed. However, ECM degradation may not be necessary for predicting gel compaction since a cluster of cells remains stationary when contracting the surrounding gel [5]. Finally, in the current work, it was assumed that the cell’s polarity mechanism is a dominating internal response to mechanical cues. Cells change their internal state through a complex process of mechanotransduction and intracellular signaling. Incorporating these more complex mechanisms is an exciting avenue for future research. While the method has been developed and demonstrated for multi-cellular interactions with 3D ECM, the basic methodology is applicable to a broad range of systems where nonlinear dynamics of many interacting subsystems are prohibitively complex to compute.

## Supporting information

**S1 Fig. Focal adhesion dynamics on an elastic substrate.** Schematic showing integrin molecules on the cellular membrane interacting with an extracellular matrix fiber, and illustrating a stochastic ligand-receptor bonding process at the focal adhesion site.

**S2 Fig. Composition of ECM fiber network model.** A: Segmented ECM fibers were generated between crosslink nodes. Yellow spheres indicate segmented ECM fiber nodes. B: A magnified view in blue circle mark in A showing examples of three fibers’ connectivity with a crosslink node. Blue lines indicated crosslinks between an ECM fiber node and a crosslink node.

**S1 Video. Comparison between original nonlinear simulation and latent variable superposition simulation of two-cell interaction embedded within cylindrical ECM.** This video depicts the cross-sectional view of the 3-D visualization of simulation of a cylindrical ECM with 2 cells embedded within it. The prediction of decreased cell volume by the latent variable superposition model (blue) agrees well with the ground-truth, full-scale nonlinear simulation results (green). The polarity directions are shown by red arrows. The polarity directions of both cells (initially pointing in arbitrary directions) shift to point inward, indicating that larger stresses are detected in the area between the cells.

**S2 Video. Comparison between two-cell latent variable superposition simulation and single cell latent variable simulation.** As can be seen from the cross-sectional view of the 3-D visualization of the simulations, the single cell model predicts more localized shrinkage of the ECM volume whereas the two cell model shows more global shrinkage extended to within the region between the cells. This suggests the presence of more than one cell is necessary for the pronounced ECM compaction leading to emergent changes within the ECM.

**S3 Video. Two-cell latent variable superposition simulation at varied spacing between 2 cells embedded within cylindrical ECM.** This video depicts the cross-sectional view of the 3-D visualization of simulation of a cylindrical ECM with 2 cells embedded within it. As the spacing between cells increases, compaction is less pronounced between them, indicating decreased interaction and integration of cell induced propagated forces.

**S4 Video. Multi-Cell latent variable superposition simulation depicting comparison of ECM compaction between heterogeneous distributions of cells.** This video depicts the cross-sectional view of the 3-D visualization of simulation of a ECM with multiple cells embedded within it. The computational model successfully reproduces the in vitro experiment conducted by Fernandez, et at [5] in which a heterogeneous planar distribution of MC3T3-E1 osteoblasts where plated in 3-D rectangular prism collagen gel. Whereas the group of 5 cells at the left edge exhibit anisotropic contraction of the ECM at the boundary, the isolated cell at the right edge does not contract the gel.

**S1 Appendix. Nonlinear dynamics of cell-ECM interaction for computational model**

**S2 Appendix. Least squares estimation for identification of the parameter matrices A, B, C, G involved in the latent space state equations**

**S3 Appendix. Implementing polarity model and lamellipodial force generation**

**S1 Table. List of simulation parameters.**

## Acknowledgments

The authors would like to thank Prof. Roger D. Kamm (MIT) and Taher Saif (UIUC) for their biological insights and advice of the studied system. The authors acknowledge support from the National Science Foundation (NSF), Science and Technology Center (STC) on Emergent Behaviors in Integrated Cellular Systems (EBICS) under Grant CBET-0939511, and Singapore-MIT Alliance of Research and Technology (SMART).

